# Emotion regulation or dual task? Dissociation of neural and behavioral measures

**DOI:** 10.64898/2026.04.17.719189

**Authors:** Nicola Sambuco, Francesco Versace, Paul M. Cinciripini, Jason D. Robinson, Yong Cui, Margaret M. Bradley, Jennifer A. Minnix

## Abstract

Cognitive reappraisal, the deliberate reinterpretation of emotional events, is widely considered an effective emotion regulation strategy, and modulation of the late positive potential (LPP) during negative affect reduction has become the primary electrophysiological evidence for volitional emotional control. Experimental instructions, however, impose dual-task demands that free viewing does not, confounding reappraisal with cognitive load. By including instructions to increase emotional responses to pictures ("enhance") as well as instructions to decrease ("suppress”), different predictions are generated. If the LPP reflects regulation, then, compared to free viewing, suppress instructions should decrease LPP amplitude, and enhance instructions should increase LPP amplitude. If modulation instead reflects cognitive load, both instructions should reduce the LPP, as both impose an additional cognitive task. In a sample of 107 participants, evaluative ratings confirmed that regulation instructions modulated reported emotional intensity in the expected directions (Enhance > View > Suppress), but that both enhance and suppress instructions reduced LPP amplitude compared to free viewing, with Bayesian model comparisons providing strong evidence against direction-specific regulation and in favor of cognitive load. Whole-scalp multivariate pattern analysis confirmed that no instruction-related neural signal exists at any scalp location or latency within the first second after stimulus onset. These data indicate that LPP modulation following both instruction types reflects dual-task cognitive load rather than volitional emotional control.

**Significance Statement:** Cognitive reappraisal is considered the gold standard of emotion regulation, and reduced late positive potential (LPP) amplitude during negative emotion suppression is the primary neural evidence that humans can voluntarily control emotional responses. The current data are inconsistent with this regulatory account and instead support a cognitive load interpretation. Whether instructed to enhance or suppress emotional responses, LPP amplitude was reduced in both conditions relative to free viewing, consistent with attentional resource competition rather than directional regulatory control. The same participants reported successfully regulating emotional experience in opposite directions, producing a clear dissociation between neural and behavioral measures. These findings challenge a basic tenet of emotional regulation and raise questions concerning LPP modulation as a biomarker of regulatory capacity.

## Introduction

An immensely popular claim in affective science is that humans can volitionally regulate their emotional responses, particularly to reduce negative affect, and that this is reflected in measurable changes in brain activity (1, 2). Among the various strategies individuals deploy to modulate their affective states, cognitive reappraisal, the deliberate reinterpretation of the meaning or personal relevance of emotional events, has received the most empirical attention (1, 3). Decades of research measuring subjective reports demonstrate that individuals instructed to “suppress” aversive stimulation using reappraisal report reduced negative affect, whereas those instructed to “enhance” pleasant experience report increased positive affect, relative to free viewing (4, 5).

At the neural level, the late positive potential (LPP) has emerged as the primary electrophysiological marker of affective processing. The LPP is a sustained centro-parietal event-related potential that is reliably enhanced by emotionally arousing stimuli relative to neutral stimuli, beginning approximately 300 ms after stimulus onset (6, 7). This enhancement is thought to reflect motivational engagement with biologically relevant information (8). Critically, LPP enhancement during picture viewing is not valence-specific. Rather, viewing either pleasant or unpleasant pictures elicits a larger LPP than viewing neutral pictures, and the magnitude of this modulation varies with motivational intensity rather than hedonic direction (9, 6, 10).

One interpretation of this pattern is that LPP amplitude indexes the allocation of processing resources to motivationally relevant stimuli (8, 9). The LPP belongs to the same family of centro-parietal positivities as the P300, a component well-established as an index of attentional resource allocation across cognitive domains (11). Consistent with a resource allocation account, when a concurrent task competes for the same limited pool of processing resources, the late positivity evoked by a primary stimulus is attenuated: P300 amplitude to target stimuli decreases systematically with increasing working memory load (12), during concurrent short-term memory scanning (13), and when a secondary task is performed during simulated driving (14).

The LPP has also been proposed as a neural index of emotion regulation success, with the majority of previous studies reporting reduced LPP when comparing free viewing of negative stimuli with free viewing under suppress instructions, interpreting this pattern as direct evidence that cognitive reappraisal reduces negative affect at the neural level (15–17). However, this design includes a critical methodological dual-task confound: during free viewing, participants attend solely to the stimulus without performing any additional cognitive operations, whereas suppress instructions require an additional, concurrent cognitive task (generating a reinterpretation). Requiring additional attentional resources is expected to divert attention from affective stimulus processing (18, 19). Because the LPP is sensitive to attentional resource allocation (9), any reduction under suppress instructions could reflect additional cognitive load rather than targeted reappraisal and reduction of the affective response.

If cognitive reappraisal genuinely alters affective processing, one should observe systematic modulation of LPP amplitude as a function of the direction of regulation instructions (15). Enhancement instructions should increase LPP amplitude relative to free viewing, whereas suppression instructions should decrease LPP amplitude. If instructions instead reflect additional cognitive demands in both dual-task conditions, LPP amplitude should be reduced in both enhance and suppress conditions, compared to free viewing. Although some studies have included instructions aimed at enhancing affective responses (20), none have crossed all three instruction conditions (free, enhance, suppress) when confronted with both pleasant and unpleasant stimuli, which provides a comprehensive assessment of how directional instructions modulate LPP amplitude.

Existing studies are often underpowered for the small-to-medium effect sizes characteristic of LPP modulation (21). In the current analysis, two independent samples of participants (total N = 107) participated in an emotion regulation task in which pleasant or unpleasant images were presented for free viewing or with instructions to enhance or suppress emotional response using cognitive reappraisal. Participants rated emotional intensity in each condition, and, replicating previous studies, ratings were expected to vary in the instructed direction (Enhance > View > Suppress). For the LPP, three competing models of effects of emotion regulation instructions were assessed using Bayesian modeling, including the regulatory reappraisal model predicting Enhance > View > Suppress, the dual-task cognitive load model predicting View > Enhance ≈ > Suppress, and a null model predicting no effects of instruction. We additionally employed whole-scalp multivariate pattern analysis to test whether instruction-related neural signals are distributed across the scalp or occur at any latency and assessed whether regulatory effects on the LPP covaried with evaluative reports of emotional intensity.

## Results

Two independent samples of participants (Sample 1, N = 73; Sample 2, N = 34) completed an emotion regulation task in which pleasant, unpleasant, and neutral pictures were first presented for free viewing. Pleasant and unpleasant pictures were then re-presented under three instruction conditions: View (continue free viewing), Enhance (use cognitive reappraisal to increase emotional response), and Suppress (use cognitive reappraisal to decrease emotional response), while high-density electroencephalography (EEG) was recorded (129 channels). Neutral scenes were only re-presented with View instructions (see **Fig. 1**) to confirm the typical pattern of increased LPP positivity for emotional, compared to neutral scenes, during free viewing.

**Figure 1.**
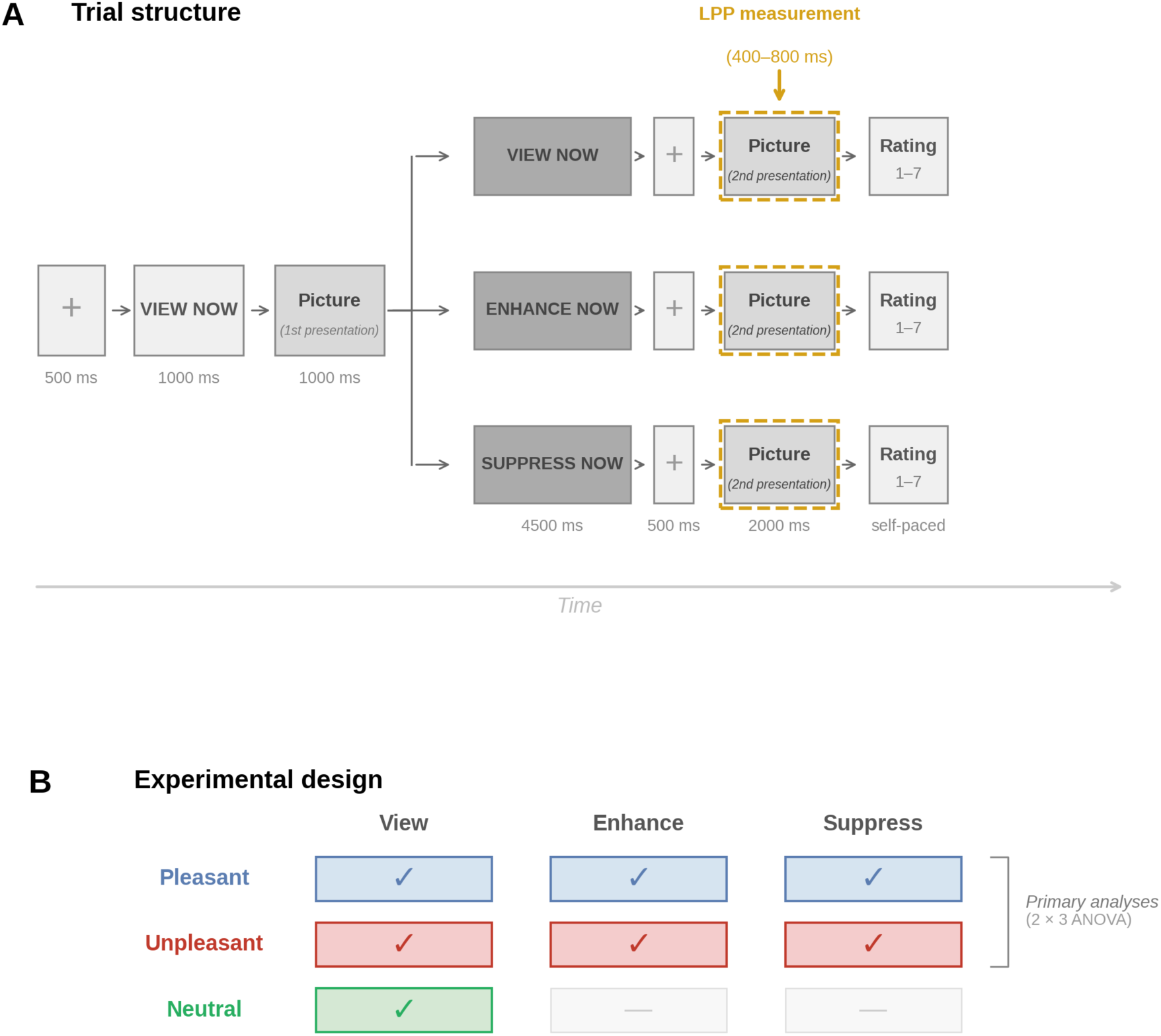
Experimental paradigm. (A) Trial structure. Each trial began with a fixation cross (500 ms) followed by an orientation cue (1000 ms) and a first picture presentation (1000 ms). Participants then received an instruction cue (4500 ms) followed by a second picture presentation (2000 ms). The LPP was measured during this second presentation (400–800 ms, dashed gold outline). (B) Experimental design. Pleasant and unpleasant pictures appeared under all three instructions (a 2 × 3 factorial design). Neutral pictures appeared only under passive viewing.

Two samples were combined to maximize statistical power. Monte Carlo simulations on LPP data collected with the same recording system used here have shown that, for within-subject effect sizes below 1 µV, achieving 80% statistical power requires at least 50 participants and that adding subjects increases power more effectively than adding trials (22). The two samples were pooled after confirming equivalent LPP responses across cohorts (see Supplementary Information).

The late positive potential (LPP) was quantified as the mean amplitude across 10 a priori centro-parietal electrode sites during the 400 to 800 ms post-stimulus window. Because participants were enrolled in a tobacco study, the stimulus set also included cigarette-related scenes, which were excluded from primary analyses (see Supplementary Results).

### Emotional intensity ratings

Emotional intensity ratings (1–7 scale) were analyzed using repeated-measures ANOVAs. During free viewing (no instructions), a one-way ANOVA confirmed that participants differentiated emotional from neutral content, F(2, 212) = 68.86, p < .001, ηG² = .278, ε = .78 (Fig. 2A). Unpleasant (M = 5.09, SD = 1.0) and pleasant (M = 4.59, SD = 1.03) pictures were rated as more emotionally intense than neutral images (M = 3.59, SD = 0.8), with unpleasant, compared to pleasant, pictures rated slightly more intense, t(106) = 3.29, p = .001, d = 0.32.

**Figure 2.**
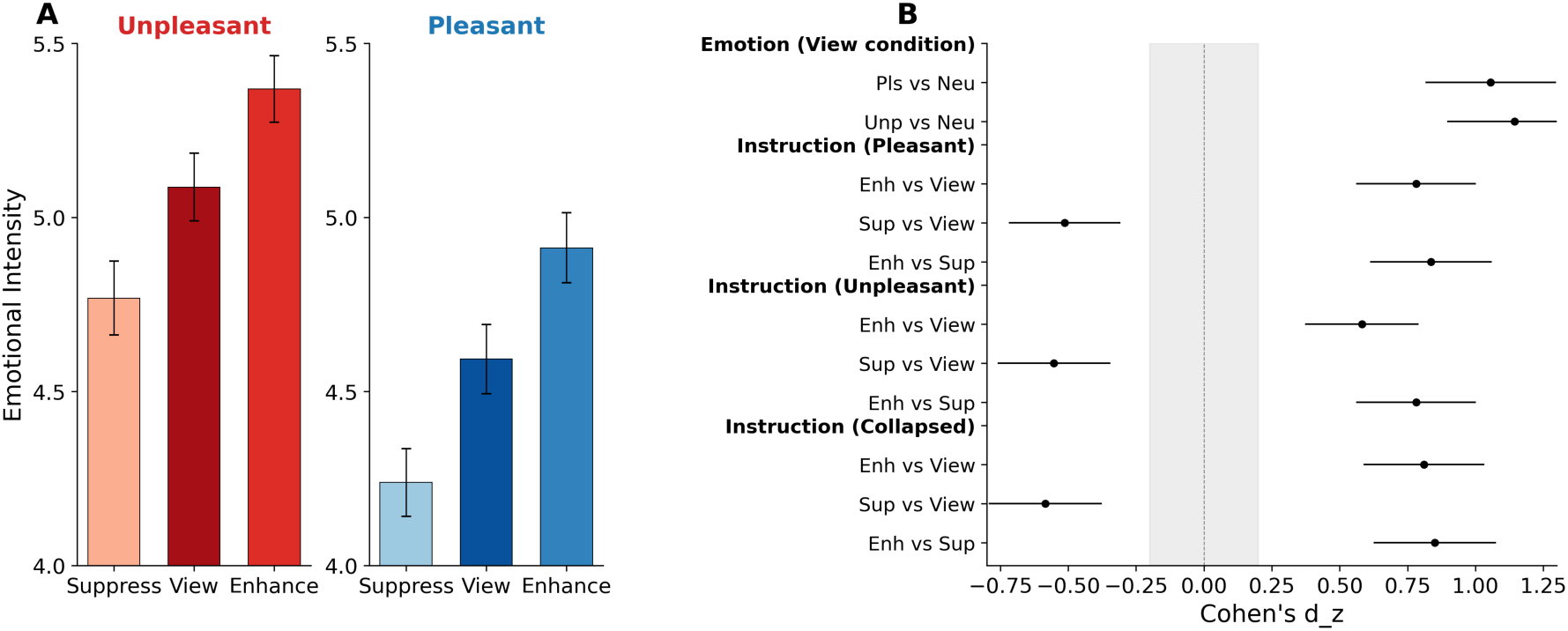
Regulation instructions modulate evaluative emotional intensity in the expected directions. (**A**) Mean ratings for pleasant (left) and unpleasant (right) pictures under Suppress, View, and Enhance instructions. Enhance instructions increased and Suppress instructions decreased intensity ratings, compared to View, for both unpleasant and pleasant scenes (all *p*s < .001). (**B**) Forest plot of within-subject effect sizes (Cohen’s *d*z) with 95% confidence intervals for all pairwise comparisons. Emotion effects in the View condition were large (*d* > 1.0); instruction effects were medium-to-large (|*d*| = 0.41–0.84). Gray shading indicates the negligible effect zone (|*d*| < 0.2). Error bars represent ±1 SEM. **p < .01, ***p < .001.

A 2 (Valence: Pleasant, Unpleasant) × 3 (Instruction: View, Enhance, Suppress) repeated-measures ANOVA for emotional pictures revealed significant main effects of Valence, F(1, 106) = 11.41, p = .001, ηG² = .050, and Instruction, F(2, 212) = 47.61, p < .001, ηG² = .057, ε = .67, with no interaction, F(2, 212) = 1.45, p = .237. Compared to free viewing, enhance instructions increased rated emotional intensity for both pleasant and unpleasant images, and Suppress instructions decreased rated emotional intensity for both as well [see **Fig. 2A**; Pleasant: Enhance (M = 4.95) > View (M = 4.60), t(106) = 7.05, p < .001, d = 0.68; View > Suppress (M = 4.27), t(106) = 4.29, p < .001, d = 0.41. Unpleasant: Enhance (M = 5.38) > View (M = 5.10), t(106) = 5.72, p < .001, d = 0.55; View > Suppress (M = 4.79), t(106) = 4.53, p < .001, d = 0.44.] Effects of instruction on evaluative reports were medium-to-large in magnitude (**Fig. 2B**). Similar effects of instruction were found when pleasant and unpleasant stimuli were divided into high and low arousal contents (see Supplementary Material).

### Affective modulation of the LPP

The initial picture presentation (View) produced the expected larger LPP for emotional compared to neutral scenes, with large effect sizes across all picture categories (see Supplementary Results).

For the second presentation, a one-way repeated-measures ANOVA under View instructions showed a significant main effect of Valence, *F*(2, 212) = 14.78, *p* < .001, η²ɢ = .058 (**Fig. 3**), with both pleasant and unpleasant scenes eliciting significantly larger LPPs than neutral scenes [Pleasant vs. Neutral, *t*(106) = 4.24, *p* < .001, *d* = 0.41 [0.21, 0.61]; Unpleasant vs. Neutral, *t*(106) = 5.10, *p* < .001, *d* = 0.49 (0.29, 0.70)]. Grand-average ERP waveforms showed clear divergence between emotional and neutral conditions beginning approximately 300 ms post-stimulus (**Fig. 3A**), and topographic maps confirmed the expected centro-parietal distribution (**Fig. 3B, C**). Similar effects were found when pleasant and unpleasant stimuli were divided into high- and low-arousal categories (see Supplementary Material).

**Figure 3.**
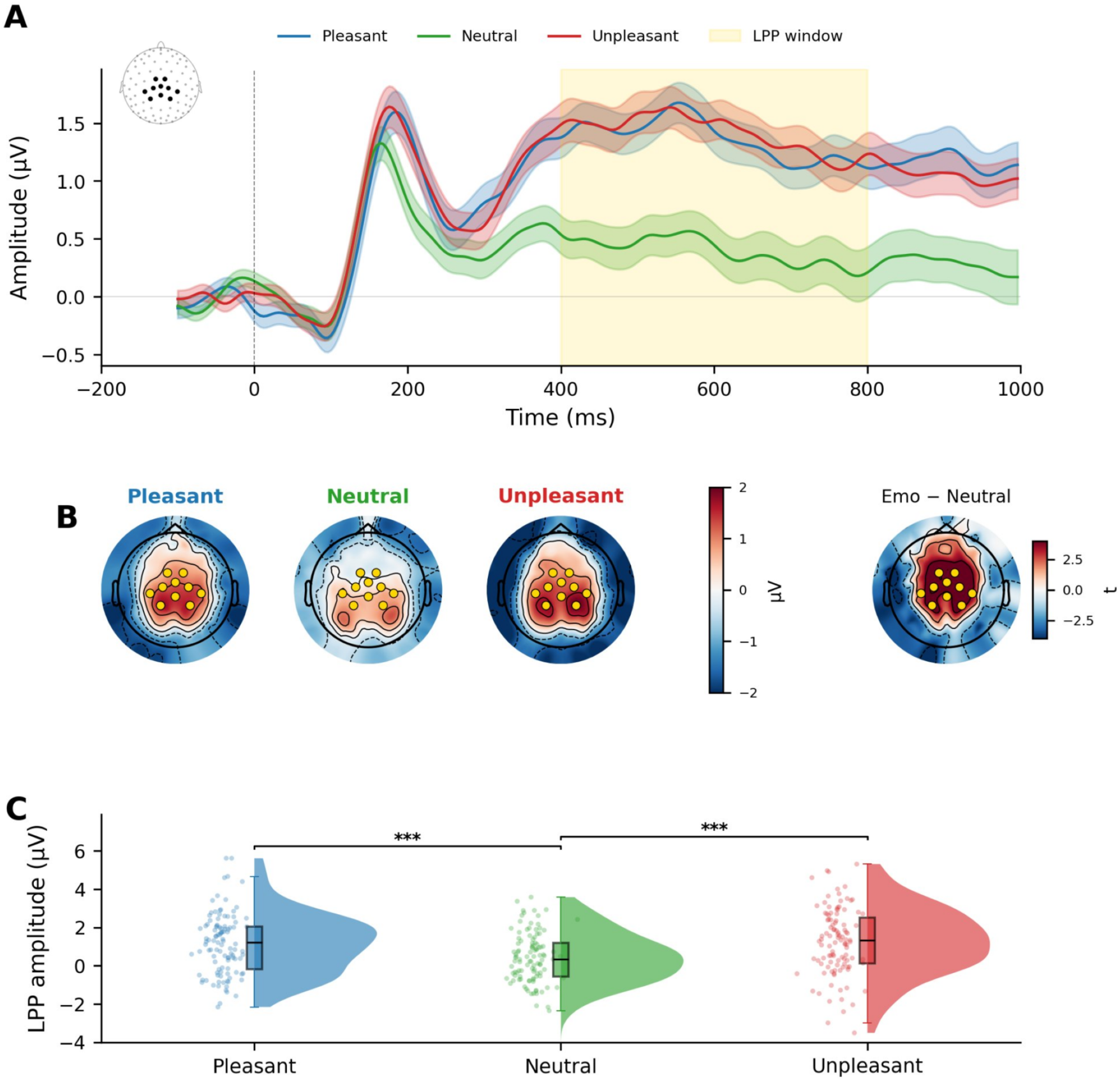
Emotional pictures elicit larger late positive potentials during free viewing. (A) Grand-average ERP waveforms at centro-parietal ROI channels (N = 107), in the inset, for pleasant (blue), neutral (green), and unpleasant (red) pictures in the View condition. Shaded ribbons represent ±SEM. Yellow shading indicates the a priori LPP measurement window (400–800 ms). (B) Topographic maps of mean scalp voltage (µV) during the LPP window. Yellow dots indicate a priori ROI channels. The rightmost map shows the Emotional − Neutral difference (t-statistic). (C) Raincloud plots of individual-participant LPP amplitudes. ***p < .001.

### Emotion regulation and the LPP

A 2 × 3 repeated-measures ANOVA with Valence (Pleasant, Unpleasant) and Instruction (Suppress, View, Enhance) was conducted on mean LPP amplitude, with a significant main effect of Instruction, F(2, 212) = 7.17, p = .001, η²ɢ = .017, but no main effect of Valence, F(1, 106) = 0.32, p = .571, and no Valence × Instruction interaction, F(2, 212) = 0.19, p = .831 (**Fig. 4A**). Critically, instead of the reappraisal-predicted ordering in which enhance instructions heightened the LPP and suppress instructions reduced the LPP, the pattern predicted by the cognitive load model was found: both enhance and suppress instructions produced reduced LPP amplitude for both pleasant and unpleasant scenes, compared to free viewing, with boths regulatory instruction eliciting similarly reduced LPPs. (**Fig. 4A**). Topographic difference maps confirmed that the View minus Enhance and View minus Suppress contrasts showed greater centro-parietal positivity, with larger LPP amplitudes found during free viewing, compared to either regulation condition (**Fig. 4B**). By contrast, the Enhance minus Suppress map showed no systematic topography, confirming that the two regulation instructions produced equivalent neural activity.

**Figure 4.**
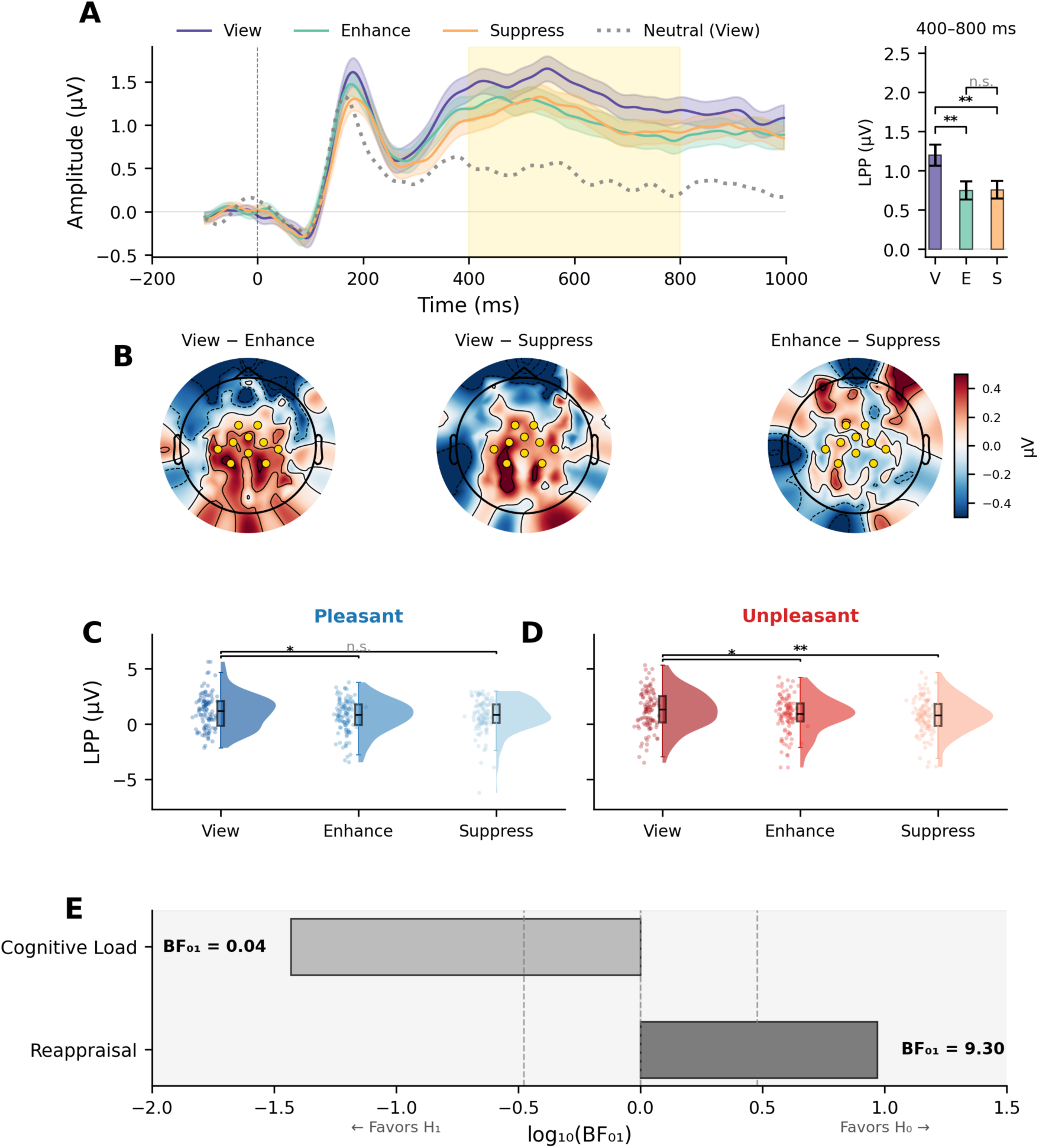
Instruction effects on the LPP are consistent with cognitive load rather than reappraisal. (**A**) Left: Grand-average ERP waveforms for emotional pictures under Suppress (amber), View (purple), and Enhance (teal) instructions. Right: Mean LPP amplitude (±SEM) in the 400–800 ms window. **p < .01, n.s. = not significant. (**B**) Topographic difference maps: View *minus* Enhance and View *minus* Suppress show centro-parietal positivity; Enhance *minus* Suppress is approximately at 0. (**C–D**) Raincloud plots of instruction effects within pleasant (**C)** and unpleasant (**D**) pictures. (**E**) Bayesian model comparison: BF₀₁ = 9.30 for the null reappraisal hypothesis; BF₀₁ = 0.04 for cognitive load.

Pairwise comparisons confirmed this pattern: For unpleasant pictures (**Fig. 4D**), LPP amplitude during View was significantly greater than during either Enhance, t(106) = 2.45, p = .016, d = 0.24 [0.04, 0.43], or Suppress instructions, t(106) = 2.69, p = .008, d = 0.26 [0.07, 0.46], which did not differ from each other. For pleasant pictures (**Fig. 4C**), LPP amplitude during View was significantly greater than during Enhance, t(106) = 2.45, p = .016, d = 0.24 [0.04, 0.43], but did not differ from Suppress, t(106) = 1.73, p = .086, d = 0.17 [−0.03, 0.36].

A Bayesian model comparison provides decisive evidence between the three hypotheses. For the Reappraisal contrast (Enhance *vs.* Suppress), the mean subject-level difference was effectively zero (MΔ = −0.01), t(106) = −0.09, p = .933, d = −0.008, BF₀₁ = 9.30, providing moderate-to-strong evidence that Enhance and Suppress produce similar LPP amplitudes (**Fig. 4E**). For the Cognitive Load contrast (2×View *minus* Enhance *minus* Suppress), the effect was robust (MΔ = 0.89), t(106) = 3.46, p < .001, d = 0.33, BF₀₁ = 0.04, indicating strong evidence for the alternative hypothesis that either regulation instruction results in significantly reduced LPP amplitude, compared to free viewing. Together, these results support a cognitive load account, rather than the pattern predicted by emotion regulation or the null model.

### Robustness analyses

Two complementary analyses confirmed the stability of the LPP findings (**Fig. 5**; see Supplementary Materials for full details). A sequential Bayes Factor analysis, progressively adding subjects from N = 20 to 107, showed that the Reappraisal contrast (Enhance vs. Suppress) consistently favored the null, with BF₀₁ rising from ≈3.5 at N = 20 to 9.30 at the full sample; the 90% credible interval remained above BF₀₁ = 1 from N ≈ 35 onward (**Fig. 5A**). The Cognitive Load contrast (View vs. Regulate) accumulated steadily in favor of the alternative, with BF₀₁ declining from ≈1.8 to 0.04 over the same range. Bootstrap effect-size distributions (10,000 resamples) converged on the same pattern (**Fig. 5B**). The Reappraisal contrast was centered on zero (d = −0.01, 95% CI [−0.20, 0.19]), whereas the Cognitive Load contrast was reliably positive (d = 0.34, 95% CI [0.17, 0.50]), with 100% of resamples exceeding zero. Neither conclusion depends on sample composition or on any small subset of participants.

**Figure 5.**
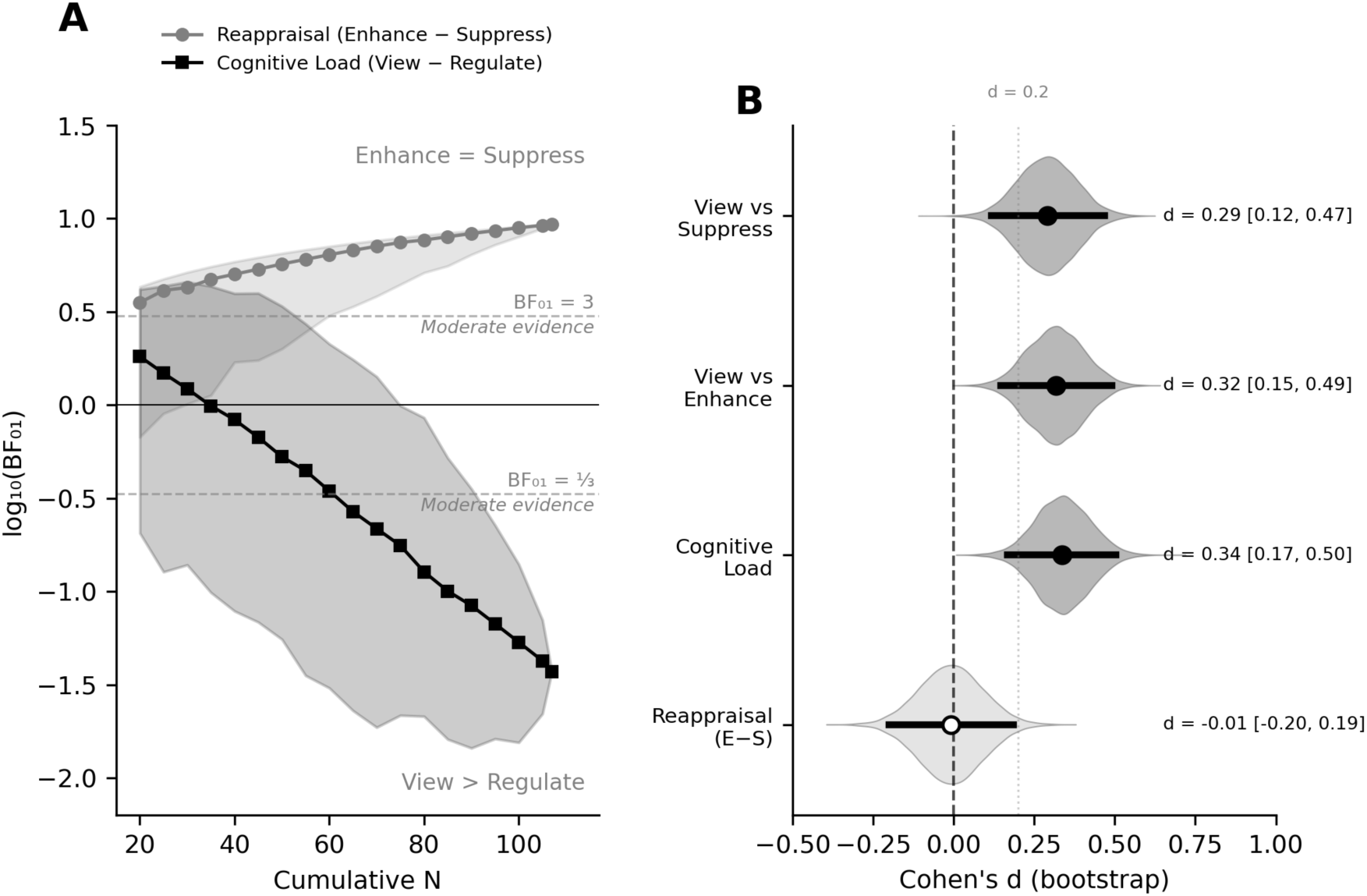
Both the reappraisal null and the cognitive load effect are robust to sample size and resampling. **(A)** Sequential Bayes factor analysis, with subjects accumulated from N = 20 to 107. The Cognitive Load contrast (black) falls steadily past the BF₀₁ = ⅓ threshold, providing increasing evidence that View exceeds both regulation conditions. The Reappraisal contrast (gray) rises above BF₀₁ = 3 by N ≈ 35, providing increasing evidence that Enhance and Suppress do not differ. Shaded bands show 90% credible intervals across 500 random subject orderings. **(B)** Bootstrap distributions of Cohen’s d (10,000 resamples). The three cognitive load comparisons (dark gray) fall entirely above zero, whereas the Reappraisal contrast (light gray) is centered on zero.

### Multivariate pattern analysis

The a priori centro-parietal ROI maximized replicability and statistical power but could, in principle, miss instruction-related activity at other scalp locations or at other latencies. To rule out this possibility, whole-scalp multivariate pattern analyses were conducted, an approach that has been used to establish multivariate neural signatures in other domains (23). Time-resolved decoding across all 129 sensors revealed that emotional content was robustly decodable. In contrast, instruction condition was not (**Fig. 6A**). Emotion decoding accuracy (Emotional vs. Neutral) exceeded the permutation threshold approximately 100 ms after stimulus onset and peaked during the LPP window (73.4%), remaining significant throughout. Instruction decoding accuracy (Enhance vs. Suppress) never exceeded the threshold at any time point.

**Figure 6.**
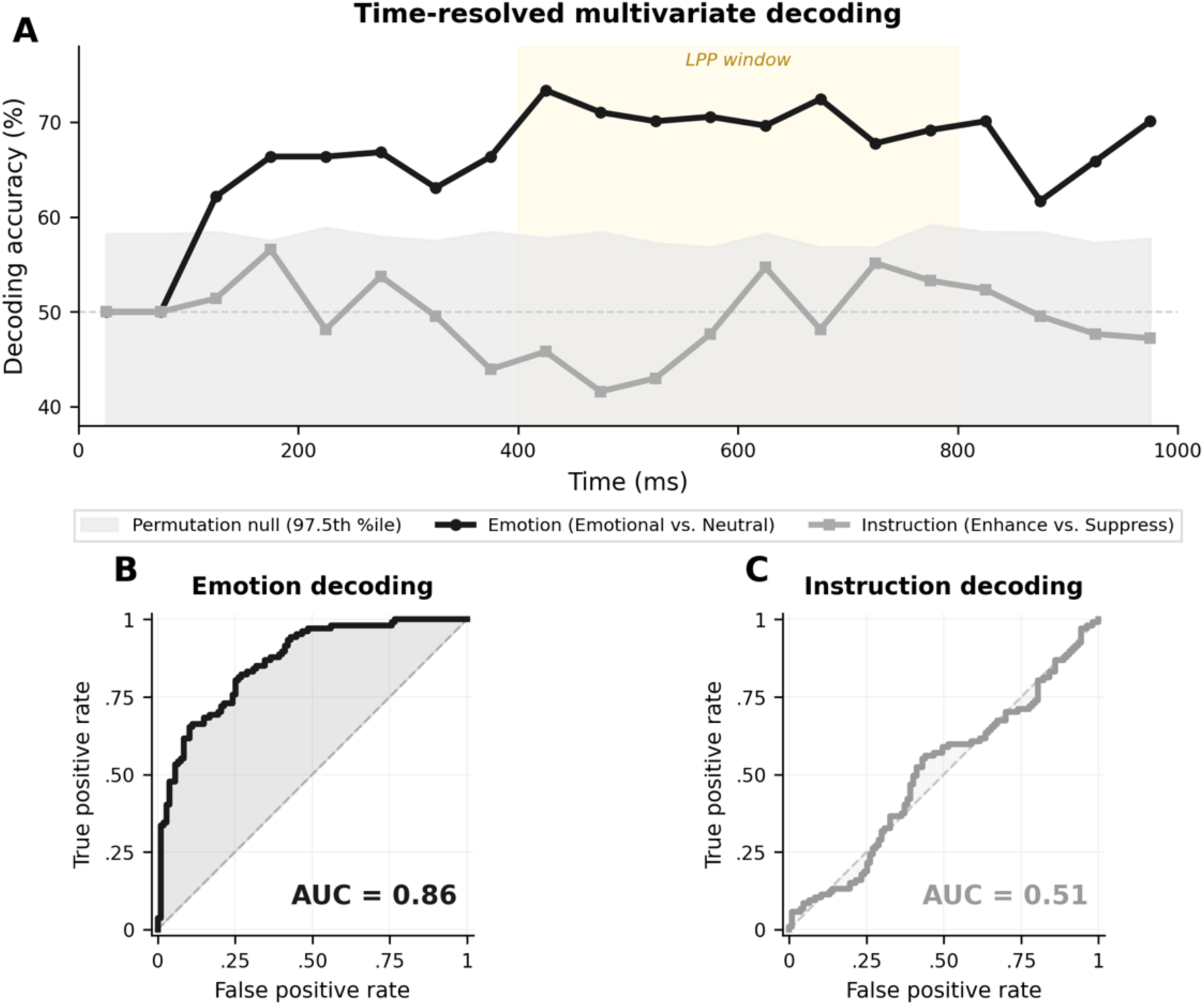
Whole-scalp decoding reads emotion from the ERP but not instruction, confirming that the reappraisal null does not reflect an insensitive ROI. (A) Time-resolved decoding accuracy across all 129 sensors. Emotional vs. Neutral classification (black) rises above the 97.5th-percentile permutation threshold (grey band) from approximately 100 ms after picture onset and peaks inside the LPP window. Enhance vs. Suppress classification (grey) stays within the permutation envelope at every time point. (B, C) ROC curves for full-epoch classification. The emotion classifier achieves AUC = 0.86, whereas the instruction classifier sits at AUC = 0.51, indistinguishable from chance. The same sensors and the same classifier that recover emotion cannot recover instruction.

Full-epoch classification confirmed this pattern (**Fig. 6B, C)**. The emotion classifier achieved 75.2% accuracy (95% CI [70.6, 79.9], AUC = 0.86, p < .001), confirming sufficient sensitivity to detect condition differences. The instruction classifier achieved 55.1% accuracy (95% CI [49.1, 61.2]), with the confidence interval including chance (AUC = 0.51, p = .077). No reliable spatiotemporal pattern distinguished Enhance from Suppress trials anywhere on the scalp or at any latency within the first second following stimulus onset.

### Brain–behavior correlations

If the LPP tracks subjective experience during emotion regulation, participants who report the largest effects of instruction on emotional experiences might show evidence of regulation-specific modulation. To test this, we correlated each participant’s rating range (Enhance − Suppress rating difference) with the corresponding LPP difference score. No relationship emerged for the pooled regulation ratings (*r* = .06, *p* = .56, BF₀₁ = 5.70; **Fig. 7A**) or the pooled suppression effect (View − Suppress; *r* = .02, *p* = .83, BF₀₁ = 6.60; **Fig. 7B**). The same null pattern held whether pleasant (*r* = .11, *p* = .24, BF₀₁ = 3.44; **Fig. 7C**) or unpleasant (*r* = .00, *p* > .99, BF₀₁ = 6.75; **Fig. 7D**) pictures were examined separately. Across all nine brain–behavior correlations (three contrasts × three valence conditions), none reached significance, and all yielded at least moderate Bayesian evidence for the null (BF₀₁ range: 3.44–6.75). Even participants who reported the strongest emotion regulation did not show LPP modulation consistent with ratings or with directional emotional regulation.

**Figure 7.**
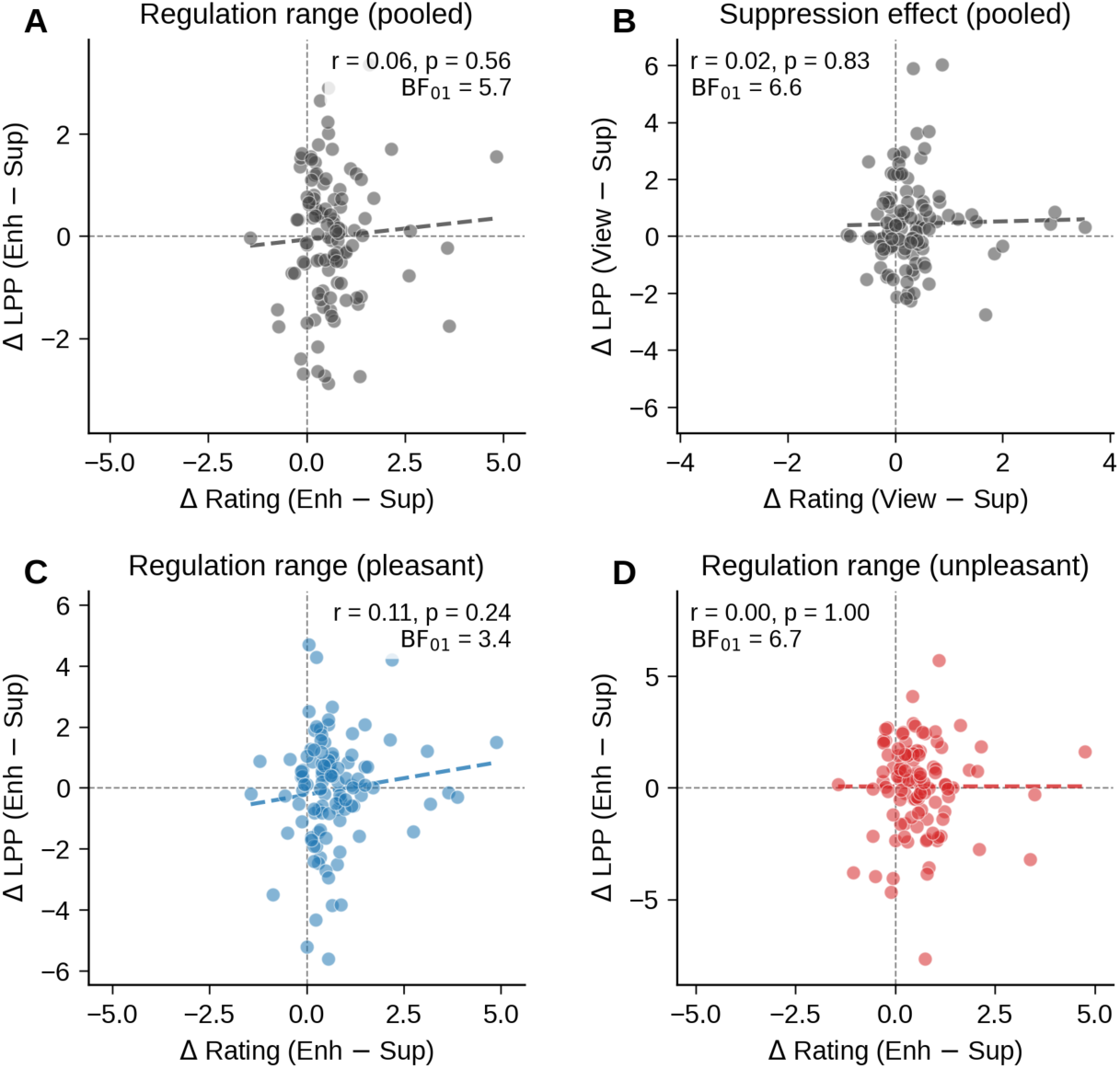
Individual differences in behavioral regulation do not predict LPP modulation. Participants who report the strongest emotion regulation show no corresponding modulation of the LPP. Each point is one participant (N = 107). The x-axis shows the magnitude of the behavioral regulation effect (difference intensity ratings), the y-axis shows the corresponding LPP difference score in µV. Dashed lines mark zero on each axis; regression lines are shown for visual reference only. (A) Pooled regulation range, Enhance − Suppress, across pleasant and unpleasant pictures (r = .06). (B) Pooled suppression effect, View − Suppress (r = .02). (C) Regulation range for pleasant pictures alone (r = .11). (D) Regulation range for unpleasant pictures alone (r = .00). All four correlations shown, and all nine brain–behavior correlations tested in total (three contrasts × three valence conditions), yield at least moderate Bayesian evidence for the null (BF₀₁ range 3.44 to 6.75). The behavioral effect of instruction and the neural effect of instruction are dissociable at the level of individual differences.

## Discussion

Participants instructed to enhance or suppress their emotional response reported emotional changes in the instructed direction, with images rated as more emotionally intense during enhance instructions and less intense during suppress instructions, compared to free viewing, replicating previous studies (4, 5, 24). At the neural level, however, the pattern was inconsistent with a regulatory account. Both enhance and suppress instructions reduced LPP amplitude when regulating unpleasant or pleasant emotion, and Bayesian model analyses provided strong evidence in favor of a cognitive load account, in which LPP amplitude varied with heightened dual-task attentional requirements, rather than direction-specific regulation of emotional responses.

Importantly, during free viewing, LPP amplitude was heightened when viewing either pleasant or unpleasant, comparewd to neutral, images, replicating decades of studies finding heightened LPP amplitude when viewing biologically relevant images (6, 9, 25). This positive control rules out poor data quality or an insensitive paradigm and confirms that, compared to free viewing, LPP amplitude was significantly reduced when either enhanced or suppressed reappraisal instructions were dual-tasks imposed during free viewing.

Whereas cognitive appraisal is considered a primary emotional regulation strategy, the same reduction in LPP amplitude during emotional picture viewing has also been found when expressive suppression, detached appraisal, or distraction are used to reduce negative affect, as shown in a recent meta-analysis of 48 LPP regulation studies (Zou et al., 2026; Qi et al, 2017). This equivalence across mechanistically distinct strategies is what a cognitive load model predicts, because all impose concurrent task demands. Consistent with this account, working memory load alone, without any regulation instruction, reduces LPP amplitude for both positive scenes (26) and during threat extinction (27), confirming that the late positivity is sensitive to concurrent cognitive demands, regardless of affective context. However, because Zou et al. ’s (2026) emotion regulation meta-analysis included only studies assessing down-regulation of negative emotion (i.e., no enhance condition), it could not distinguish between models. A previous study examining only upregulation for pleasant (and neutral) scenes reported main effects of valence and task on LPP amplitude, but no valence-by-instruction interaction that would support direction-specific upregulation of positive emotion (20). The present study fills the gap by measuring ERPs when participants were instructed to enhance negative, as well as positive emotional experience, with regulation and cognitive load models generating divergent predictions. The results are unambiguous: all dual-task instructional conditions reduced LPP amplitude, compared to free viewing.

Whereas a reduction in attention to the primary picture viewing task by imposing a secondary, concurrent task may mirror the desired regulation effect for "suppress" instructions during aversive image processing (e.g. reduce negative image processing), it will not mirror desired effects when enhancing emotional response to negative stimuli (which is rarely desirable in clinical contexts) or when enhancing emotional response to positive emotion, which is often the clinical goal (28, 29). A demanding secondary task, such as cognitive reappraisal, which requires participants to generate additional complex scenarios, instead seems to require substantial resources, reducing the amplitude of the LPP found during emotional picture viewing for both positive and negative scenes, whether instructed to enhance or suppress emotional experience (30).

An important question is whether the present findings extend beyond the LPP to other neural measures of emotion regulation. As for the LPP, the bulk of functional MRI studies assess neural activity during the downregulation of negative emotion (31). Meta-analyses of neural activation report increased prefrontal cortex activation and reduced amygdala activity, which are similar to regions identified as active in demanding dual-task situations. For instance, a meta-analysis of the 2-back working memory task (32) also reports enhanced prefrontal activation and bilateral amygdala deactivation clusters that overlap substantially with those observed during suppress cognitive reappraisal.

A central claim of emotion regulation research is that activation of fronto-parietal networks exerts volitional, direction-specific control over subcortical emotion-generative systems (33), functioning as a bidirectional gain mechanism that can attenuate or amplify affective responses on command (2, 34–37). Enhance conditions, which are the least investigated in previous studies, comprise the strongest test of this bidirectional assumption but the few neuroimaging studies that have included upregulation instructions have used small samples with limited statistical power, and meta-analyses that include these data result in small, scattered prefrontal and subcortical activations that do not converge on a robust or replicable pattern (31). High-powered within-subject studies that manipulate both enhance and suppress for both pleasant and unpleasant stimuli are needed to determine whether fronto-parietal activation varies directionally with instruction or instead reflects nonspecific cognitive demands common to all regulation conditions.

Although the LPP was reduced following both enhance and suppress instructions, participants reported experiencing greater emotional intensity following enhance reappraisal and reduced emotional intensity following suppress reappraisal, compared to free viewing, consistent with many previous studies reporting that regulation instructions modulate evaluative reports in the experimenter-requested direction (5, 16). Nonetheless, although reports of emotional experience covaried with instructed direction of emotional change, the neural LPP index only showed significant attenuation under both regulation instructions. Even participants who reported the strongest emotion regulation showed no corresponding differential LPP modulation. Regulation instructions achieve the instructed goal for reports of emotional experience but do not produce corresponding directional changes in the LPP. This is consistent with a recent large-scale consortium study (38), which found that relationships between ratings following emotion regulation and neural activation were extremely small or nonexistent in analyses that included thousands of participants, and with broader evidence that subjective emotional arousal and LPP amplitude dissociate for several stimulus categories (39).

This raises a critical issue of the extent to which modulatory effects of regulation instructions on reports of emotional experience are primarily mediated by demand effects (40), in which participants understand the goal of the study and adjust their responses accordingly. Demand is particularly suspect in emotion regulation tasks, as instructions explicitly ask participants to increase or decrease emotional intensity, clearly communicating the experimental goal (41). In the absence of any effort to disguise the critical experimental hypotheses, dependent measures that are volitionally controlled, such as reports of affective experience, are highly susceptible to demand characteristics. In this case, other measures, such as the electrophysiological LPP, are needed to provide evidence that demand is not mediating subjective reports; yet the LPP did not vary with the direction of the regulation instruction or covary with the degree of reported regulation success, making a demand interpretation of evaluative reports possible.

Reappraisal-based interventions are among the most widely used treatments for emotional dysregulation, and LPP modulation has been proposed as a biomarker of regulatory success (15, 42). To the extent that emotion regulation strategies such as cognitive reappraisal and distraction both divert processing resources away from aversive stimulation (43), little is added to the original coping framework, in which numerous strategies have long been recognized as effective in reducing aversive processing (44). More importantly, the current data suggest that instructions to enhance emotional experience also attenuate LPP amplitude, due to dual-task requirements, rather than signaling heightened emotional engagement. This is a critical issue in recent efforts to harness emotion regulation to upregulate blunted positive affect in depression and anhedonia, which rely primarily on evaluative reports to index regulatory success or capacity (45).

In summary, in a pooled sample of 107 participants with Bayesian model comparison, the LPP in the 400–800 ms window was sensitive to the motivational relevance of visual images during free viewing and was reduced by the cognitive demands imposed by concurrent emotion regulation instructions, regardless of whether the instructed direction was to suppress or enhance negative or positive reactions. Replicating many previous studies, participants reported directional changes in the intensity of emotional experience consistent with regulation instructions, however reports did not covary with neural responses. Because emotion dysregulation has been proposed to mediate a large variety of mental health disorders, including anxiety, depression, eating and substance use disorders, borderline personality disorder, etc. (46, 47), it is critical to determine if any neural measures covary with reports of emotional experience, as predicted by an emotion regulation account. This is especially relevant given growing efforts to develop transdiagnostic neural indicators of affective dysfunction in clinical research (48) that can assist in diagnosing and treating mental health disorders.

## Methods

### Participants

Participants were adult daily smokers recruited from the Houston (TX) community who reported smoking at least 10 cigarettes per day for at least one year. Exclusion criteria included current psychiatric or neurological disorders, use of psychotropic medication, and contraindications for EEG recording. Data were collected from two independent samples using the same paradigm and equipment. Sample 1 comprised N = 73 participants after exclusions for excessive EEG artifacts; Sample 2 comprised N = 34 participants. Both samples were drawn from the same community and completed identical procedures. All participants provided written informed consent, and the protocol was approved by the local Institutional Review Board.

The pooled sample (N = 107) included 61 female and 46 male participants (M age = 45.6 years, SD = 10.2, range: 21–68). Participants self-identified as White (56.1%), Black or African American (39.3%), more than one race (1.9%), or Hispanic or Latino (0.9%); 1.9% preferred not to disclose their race. Eight participants (7.5%) identified as Hispanic or Latino ethnicity. Sample 1 (n = 73; 41 female, 32 male; M age = 48.0 years, SD = 10.1, range: 25–68) was predominantly White (71.2%) and Black or African American (24.7%), with 9.6% identifying as Hispanic or Latino. Sample 2 (n = 34; 20 female, 14 male; M age = 40.4 years, SD = 8.4, range: 21–50) was predominantly Black or African American (70.6%) and White (23.5%); ethnicity data were available for 18 of 34 participants, of whom one (5.6%) identified as Hispanic or Latino.

### Task and stimuli

The cognitive reappraisal task (**Fig. 1**) was built using E-Prime. Stimuli were affective pictures drawn from the IAPS (49) and other sets, organized into six categories: pleasant high-arousal, pleasant low-arousal, unpleasant high-arousal, unpleasant low-arousal, neutral, and cigarette-related. Each trial consisted of two successive presentations of the same picture. During the first presentation, all pictures were passively viewed. Participants then received an instruction cue (VIEW, ENHANCE, or SUPPRESS) followed by the same picture a second time. In the View condition, participants continued to attend naturally. In the Enhance condition, participants used cognitive reappraisal to increase their emotional response, for example by focusing on a specific feature of the picture or imagining an outcome that would intensify the emotion. In the Suppress condition, participants used cognitive reappraisal to decrease their emotional response, for example by focusing on a positive aspect of the picture or imagining a positive outcome of the depicted situation.

Participants were explicitly instructed not to generate thoughts unrelated to the picture (see Supplementary Methods for the exact instruction wording). Following the second presentation, participants rated the emotional intensity of their response on a 1 to 7 scale. Neutral pictures appeared only under View. Prior to the task, participants received reappraisal training and completed practice trials. The task was divided into two halves of four blocks each, separated by an electrode impedance check. Instruction condition was distributed equally within each block, and specific picture-instruction pairings were counterbalanced across participants using predefined lists.

### EEG recording and processing

Continuous EEG was recorded using a 129-channel HydroCel Geodesic Sensor Net (EGI) connected to a Net Amps 300 amplifier at 250 Hz, with Cz as the online reference. Electrode impedances were below 50 kΩ. Offline, data were bandpass filtered and segmented into epochs time-locked to picture onset. Epochs were baseline-corrected and subjected to automated artifact rejection. Bad channels were interpolated using spherical spline interpolation. Data were re-referenced to the average reference (see Supplementary Methods for full details).

### Statistical analysis

All analyses were performed using Python 3.10 (pingouin 0.5, scipy 1.11). Emotional intensity ratings (1–7 scale) were averaged within each Valence × Instruction cell for each participant and analyzed using repeated-measures ANOVAs with Greenhouse-Geisser correction. Within-subject effect sizes (Cohen’s d*z*) were computed with 95% confidence intervals. The LPP was quantified as the mean amplitude across 10 a priori centro-parietal electrode sites (EGI channels 7, 31, 37, 54, 55, 79, 80, 87, 106, and 129) during the 400–800 ms window, selected based on previous work (22, 50). LPP values were winsorized at the 1st and 99th percentiles. Data from both samples were pooled (N = 107) after confirming that the samples did not differ in LPP responses (see Supplementary Materials). To distinguish between reappraisal and cognitive load accounts, we tested two orthogonal contrasts: (1) Reappraisal (Enhance − Suppress), and (2) Cognitive Load (2×View − Enhance − Suppress). For each, we computed Bayes Factors (BF₀₁) using the JZS prior (Cauchy scale r = √2/2) (51). BF₀₁ values between 3 and 10 indicate moderate evidence for the null; values above 10, strong evidence. Values between 1/3 and 1/10 indicate moderate evidence for the alternative; below 1/10, strong evidence (52). Full details of all statistical procedures, including robustness analyses, arousal analyses, and MVPA pipeline, are provided in Supplementary Methods. To assess brain–behavior relationships at the individual level, we correlated each participant’s behavioral regulation score (Enhance − Suppress rating difference) with the corresponding LPP difference score, separately for pleasant, unpleasant, and pooled conditions. Pearson correlations were accompanied by JZS Bayes factors (prior scale κ = √2/2) (51).

## Acknowledgments and funding sources

This work was supported by National Institutes of Health grants K99DA025181 and R00DA025181 (to J.A.M.) and R01DA024709 (to P.M.C.) from the National Institute on Drug Abuse, and by National Cancer Institute Cancer Center Support Grant P30CA016672 to The University of Texas MD Anderson Cancer Center.

## Data, Materials, and Software Availability

Individual-level EEG and behavioral data are not publicly shared because the approved study protocol and participant consent procedures do not permit public release of human subjects data. De-identified data supporting the findings reported here can be made available to qualified researchers upon request from one of the corresponding authors, under a data use agreement consistent with the approved IRB protocol. All analysis code is publicly available at https://github.com/nsambuco/LPP_reappraisal, including statistical analysis scripts and figure generation code. The repository includes demonstration scripts with example inputs that reproduce the analytical workflow reported in the main text and Supplementary Materials.

## SUPPLEMENTARY INFORMATION

### Supplementary Methods

#### Regulation instruction wording

Participants were trained on three instruction conditions prior to the task. The following wording was used:

##### View

"Simply look at the picture."

##### Enhance

"We would like you to increase the intensity of the emotion you feel in response to the picture. Try to feel the emotion more strongly. Do not generate thoughts and images that are completely unrelated to the picture in order to produce a different emotion. However, you may focus on a specific feature of the picture or think of an outcome of the situation depicted in the picture that will help to enhance your emotion. Prepare yourself to feel the emotion more strongly."

##### Suppress

"We would like you to decrease the intensity of the emotion you feel in response to the picture. Try to feel the emotion less strongly. Suppression of an emotion is not the same as replacing that emotion with a different one. Do not generate thoughts and images that are completely unrelated to the picture in order to produce a different emotion. However, feel free to focus on a positive aspect of the picture or on a possible positive outcome of the situation in the picture. Prepare yourself to feel the emotion less strongly."

Participants practiced each instruction with feedback from the experimenter until they could provide at least two adequate examples of each strategy

#### Rationale for combining samples

Given the small-to-medium effect sizes typically reported for cognitive reappraisal modulations of the LPP (Zou et al., 2026), the two samples were combined to maximize statistical power. Monte Carlo simulations on LPP data collected with the same recording system used here have shown that, for within-subject effect sizes below 1 µV, achieving 80% statistical power requires at least 50 participants and that adding subjects increases power more effectively than adding trials (Gibney et al., 2020). Because the instruction effects of interest fall well below 1 µV, neither Sample 1 (N = 73) nor Sample 2 (N = 34) alone provides adequate power, particularly for the critical Enhance versus Suppress contrast. Combining was contingent on two prerequisites: (a) a sensitivity power analysis confirming inadequate individual power, and (b) a mixed-model ANOVA confirming equivalent LPP responses across samples. Both conditions were met (see below).

#### EEG preprocessing details

Offline, continuous EEG data were visually inspected, and broken channels were interpolated using spherical splines. After re-referencing to the average of all electrodes, eye blinks were corrected using a spatiotemporal filtering method as implemented in BESA 7.1.2.1 (BESA GmbH, Gräfelfing, Germany). Subsequent analyses were performed in BrainVision Analyzer 2.3.0 (Brain Products GmbH, Gilching, Germany). Data were bandpass-filtered to 0.1-40 Hz and segmented into 1100 ms epochs time-locked to picture onset. Epochs were baseline-corrected using the 100 ms pre-stimulus interval. Automated artifact detection was applied to each channel to identify epochs contaminated by artifacts. The following criteria defined artifacts: EEG amplitude above 100 or below -100 μV during the epoch; absolute voltage difference between any two data points within the segment larger than 100 μV; voltage difference between two contiguous data points above 25 mV and less than 0.5 μV variation for more than 100 ms. Channels contaminated by artifacts in more than 40% of the epochs were interpolated using spherical splines. Epochs with more than 10% of channels contaminated by artifacts were discarded. After these steps, ERPs were calculated for each channel and each condition.

#### Detailed statistical procedures

##### Emotional intensity ratings

Trial-level emotional intensity ratings (1–7 scale) were averaged within each Valence × Instruction cell for each participant. A 2 (Valence: Pleasant, Unpleasant) × 3 (Instruction: View, Enhance, Suppress) repeated-measures ANOVA tested whether regulation instructions modulated subjective experience. The emotion effect was tested separately with a one-way RM ANOVA on View-condition ratings (Neutral, Pleasant, Unpleasant). All ANOVAs used Greenhouse-Geisser correction where sphericity was violated. Follow-up pairwise comparisons used paired t-tests with Cohen’s d as the within-subject effect size. Cigarette-related pictures were analyzed separately.

##### LPP: Emotion effect

To confirm LPP sensitivity to emotional content, we tested the effect of picture valence in the View condition only using a one-way repeated-measures ANOVA with Valence (Pleasant, Neutral, Unpleasant). The Greenhouse-Geisser correction was applied when sphericity was violated. Follow-up comparisons contrasted each emotional category against Neutral using two-tailed paired t-tests. Effect sizes are reported as Cohen’s d with 95% confidence intervals.

##### LPP: Reappraisal model testing

A 2 × 3 repeated-measures ANOVA with Valence (Pleasant, Unpleasant) and Instruction (View, Enhance, Suppress) was conducted. The standard paradigm introduces a confound: View is a single-task condition, whereas Enhance and Suppress are dual-task conditions. To adjudicate between accounts, we adopted a Bayesian model-comparison framework testing two orthogonal contrasts:

(1) Reappraisal model (Enhance − Suppress, [0, +1, −1]). If participants can volitionally regulate the LPP, Enhance should exceed Suppress. (2) Cognitive load model (2×View − Enhance − Suppress, [+2, −1, −1]). If both regulation instructions divert attentional resources, both should reduce the LPP equally relative to View. These contrasts are mathematically orthogonal and provide independent tests. For each, we computed a one-sample t-test against zero on subject-level contrast scores (collapsed across valences) and derived BF₀₁ using the JZS prior (Cauchy scale r = √2/2 ≈ 0.707; Rouder et al., 2009) as implemented in the pingouin library.

##### LPP: Robustness analyses

Two complementary checks were performed. First, a sequential Bayes Factor analysis evaluated evidence accumulation across participants. Subjects were randomly ordered (500 permutations), and BF₀₁ for both contrasts was computed at incremental sample sizes (N = 20, 25, …, 107). Median BF₀₁ values and 90% credible intervals are reported. Second, a nonparametric bootstrap analysis (10,000 resamples) estimated the sampling distribution of Cohen’s d for each contrast.

##### LPP: Arousal × Instruction analysis

We examined whether instruction effects varied with stimulus arousal. The original stimulus set included high-arousal and low-arousal subcategories within each valence. We compared View to a collapsed Regulate condition (mean of Enhance and Suppress) separately for high- and low-arousal pictures. Bayesian t-tests were computed for both the Reappraisal and View vs. Regulate contrasts at each arousal level.

##### Multivariate pattern analysis

To test whether instruction-related signals are present anywhere on the scalp, we applied multivariate pattern analysis (MVPA) using a two-step positive-control approach. The same pipeline was used for: (1) emotion decoding (Emotional vs. Neutral, View condition; positive control), and (2) instruction decoding (Enhance vs. Suppress, emotional pictures; critical test). For emotion classification, the Emotional class was the average of Pleasant-High and Unpleasant-High conditions under View; Neutral was the Neutral-View condition. For instruction classification, Enhance was the average of Pleasant-Enhance and Unpleasant-Enhance; Suppress was the average of Pleasant-Suppress and Unpleasant-Suppress. Each subject contributed one observation per class.

Feature vectors comprised mean voltage at all 129 channels within each of 20 consecutive 50-ms time bins spanning 0–1000 ms, yielding 2,580 features per observation. The pipeline consisted of z-score standardization, PCA retaining 95% of variance, and L2-regularized logistic regression (C = 1.0), with all transformations fit on training data only within each cross-validation fold. Leave-one-subject-out cross-validation (107 folds) was used. Statistical significance was assessed by permutation testing (1,000 iterations). Time-resolved analysis trained separate classifiers at each 50 ms bin using standardization and logistic regression without PCA, with permutation testing (100 iterations per bin).

## Supplementary Results

### Sensitivity Power Analysis

To evaluate whether pooling the two samples was warranted, we conducted a sensitivity power analysis for paired *t*-tests across a range of Cohen’s *d* values (0.05 to 0.80) and sample sizes (10 to 300), with alpha = .05 (two-tailed). Power was computed analytically using the non-central *t* distribution. The observed effect sizes from the pooled sample were then projected onto these curves to determine the power available at each sample size (Supplementary Figure 1).

For the basic emotion effect (emotional vs. neutral pictures under passive viewing), the observed effect sizes were medium in magnitude (*d* = 0.41 for pleasant, *d* = 0.49 for unpleasant). Sample 1 (*N* = 73) was adequately powered for both contrasts (power = .93 and .99, respectively), whereas Sample 2 (*N* = 34) reached 80% power only for the unpleasant contrast. The critical effects for the present study, however, are the instruction contrasts. The observed cognitive load effect (View vs. Regulation) yielded *d* = 0.257, and the individual pairwise contrasts (View vs. Enhance, View vs. Suppress) ranged from *d* = 0.236 to 0.261. At these effect sizes, Sample 2 afforded only 27-32% power and Sample 1 only 51-59% power, both well below the conventional 80% threshold. Achieving 80% power for these contrasts requires between 118 and 143 participants, confirming the need for pooling. Notably, the Enhance versus Suppress contrast, the key test of the reappraisal account, yielded a near-zero observed effect (*d* = 0.030), which would require 80% power with over 8,700 participants, reinforcing the conclusion that the absence of a reappraisal effect is not attributable to insufficient sample size. These results converge with Monte Carlo simulations conducted on LPP data collected with the same recording system, which demonstrated that within-subject effects below 1 µV require at least 50 participants to reach adequate power.

**Supplementary Figure 1.**
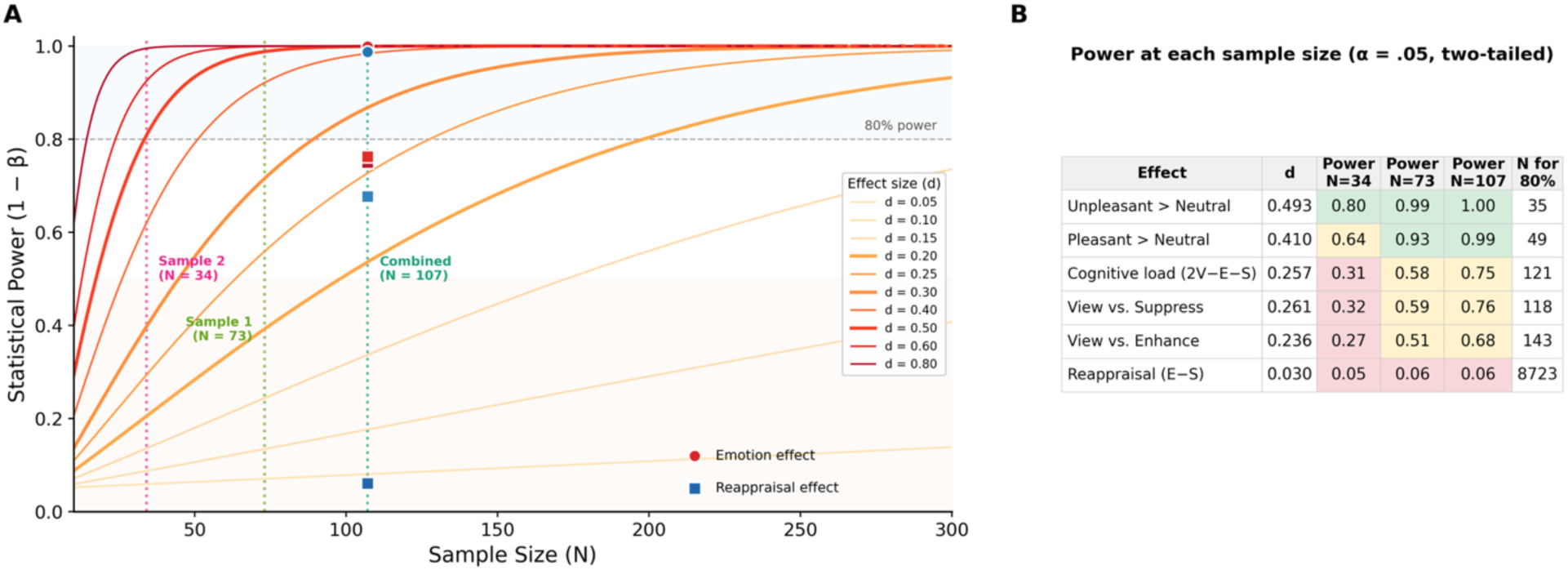
Sensitivity power analysis justifying sample pooling. (A) Statistical power (1 − β) as a function of sample size for paired *t*-tests at effect sizes ranging from *d* = 0.05 to *d* = 0.80 (α = .05, two-tailed). Vertical dotted lines indicate Sample 2 (*N* = 34), Sample 1 (*N* = 73), and the combined sample (*N* = 107). Colored markers show observed effect sizes from the pooled analysis projected onto the combined sample size: circles denote emotion effects (emotional vs. neutral, View condition) and squares denote instruction effects. The dashed horizontal line marks 80% power. (B) Power estimates for each observed effect at each sample size, with the sample size required to achieve 80% power. Green cells indicate adequate power (≥ .80), yellow cells marginal power (.50 to .79), and red cells insufficient power (< .50).

### Cohort Equivalence

To confirm that the two samples produced comparable LPP responses, we conducted a mixed-model ANOVA on LPP amplitudes in the View condition, with Valence (Pleasant, Unpleasant, Neutral, Cigarette) as a within-subjects factor and Cohort (Sample 1, Sample 2) as a between-subjects factor. The two samples did not differ in overall LPP amplitude, *F*(1, 105) = 2.61, *p* = .109, η²p = .024, and the Valence × Cohort interaction was not significant, *F*(3, 315) = 0.52, *p* = .667, η²p = .005, indicating that the pattern of valence-dependent LPP modulation was consistent across samples. The main effect of Valence was significant, *F*(3, 315) = 12.58, *p* < .001, η²p = .107, confirming that the basic emotion effect replicated in both samples. Having satisfied both prerequisites for pooling (insufficient power in individual samples and equivalent LPP responses across cohorts), all subsequent analyses were conducted on the combined sample (*N* = 107).

### Regulation Effect in Self-Report Ratings

To test whether behavioral regulation varied with stimulus intensity, we conducted separate 2 (Arousal: High, Low) × 3 (Instruction: View, Enhance, Suppress) repeated-measures ANOVAs for pleasant and unpleasant pictures (Supplementary Fig. 2). For pleasant pictures, both main effects were significant, Arousal, *F*(1, 105) = 96.69, *p* < .001, η²G = .081, and Instruction, *F*(2, 210) = 56.84, *p* < .001, η²G = .062, ε = .70, with no Arousal × Instruction interaction, *F*(2, 210) = 2.18, *p* = .115, indicating that regulation effects were comparable across arousal levels. For unpleasant pictures, both main effects were again significant, Arousal, *F*(1, 105) = 103.38, *p* < .001, η²G = .112, and Instruction, *F*(2, 210) = 49.12, *p* < .001, η²G = .046, ε = .77, and the interaction reached significance, *F*(2, 210) = 5.92, *p* = .003, reflecting a smaller enhancement effect for high-arousal (*d* = 0.32) than low-arousal (*d* = 0.67) unpleasant pictures. Critically, all 12 pairwise contrasts were significant (all *p*s < .01, |*d*| = 0.32–0.82; **Fig. 2F**), confirming that behavioral regulation was effective at both arousal levels for both valence categories.

**Supplementary Figure 2.**
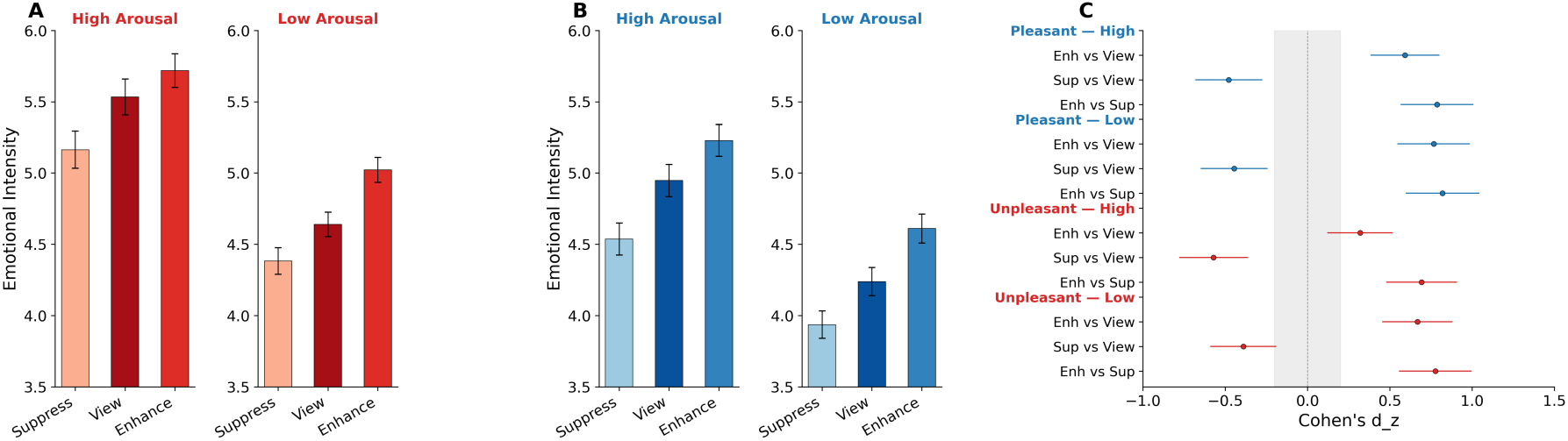
(**A**) Mean ratings for pleasant pictures by arousal level (High, Low) and instruction condition. (**B**) Mean ratings for unpleasant pictures by arousal level and instruction condition. Regulation effects were significant at both arousal levels for both valence categories (all *p*s < .01). (**C**) Forest plot of arousal-specific within-subject effect sizes. **p < .01, ***p < .001.

### First View: Emotion Effect Prior to Instruction

To verify that the LPP was sensitive to emotional content before any regulatory instructions were introduced, we examined the first picture presentation, during which all pictures were passively viewed regardless of the upcoming instruction condition. Grand-average ERP waveforms showed clear differentiation among all four valence categories beginning approximately 200 ms post-stimulus, with pleasant and unpleasant pictures producing the largest sustained positivity across the LPP window (Supplementary Fig. 3A). Cigarette pictures tracked between emotional and neutral conditions throughout the epoch. Topographic maps of the Emotional − Neutral difference confirmed a centro-parietal distribution emerging by 200–400 ms and strengthening through the LPP window (Supplementary Fig. 2B, top row), and individual valence topomaps at 400–800 ms showed the expected pattern of broadly distributed positivity for all stimulus categories relative to neutral (Supplementary Fig. 3B, bottom row).

A one-way repeated-measures ANOVA on mean LPP amplitude (400–800 ms) with Valence (Pleasant, Neutral, Unpleasant, Cigarette) confirmed a significant main effect, *F*(3, 318) = 71.39, *p* < .001, η²G = .123, ε = .86 (Supplementary Fig. 3C). All three stimulus categories elicited significantly larger LPPs than Neutral: Pleasant, *t*(106) = 12.87, *p* < .001, *d* = 1.24 [1.00, 1.50]; Unpleasant, *t*(105) = 9.89, *p* < .001, *d* = 0.96 [0.73, 1.19]; and Cigarette, *t*(106) = 5.76, *p* < .001, *d* = 0.56 [0.35, 0.76]. Pleasant pictures also elicited larger LPPs than Unpleasant pictures, *t*(105) = 6.39, *p* < .001, *d* = 0.62 [0.41, 0.83].

**Supplementary Figure 3.**
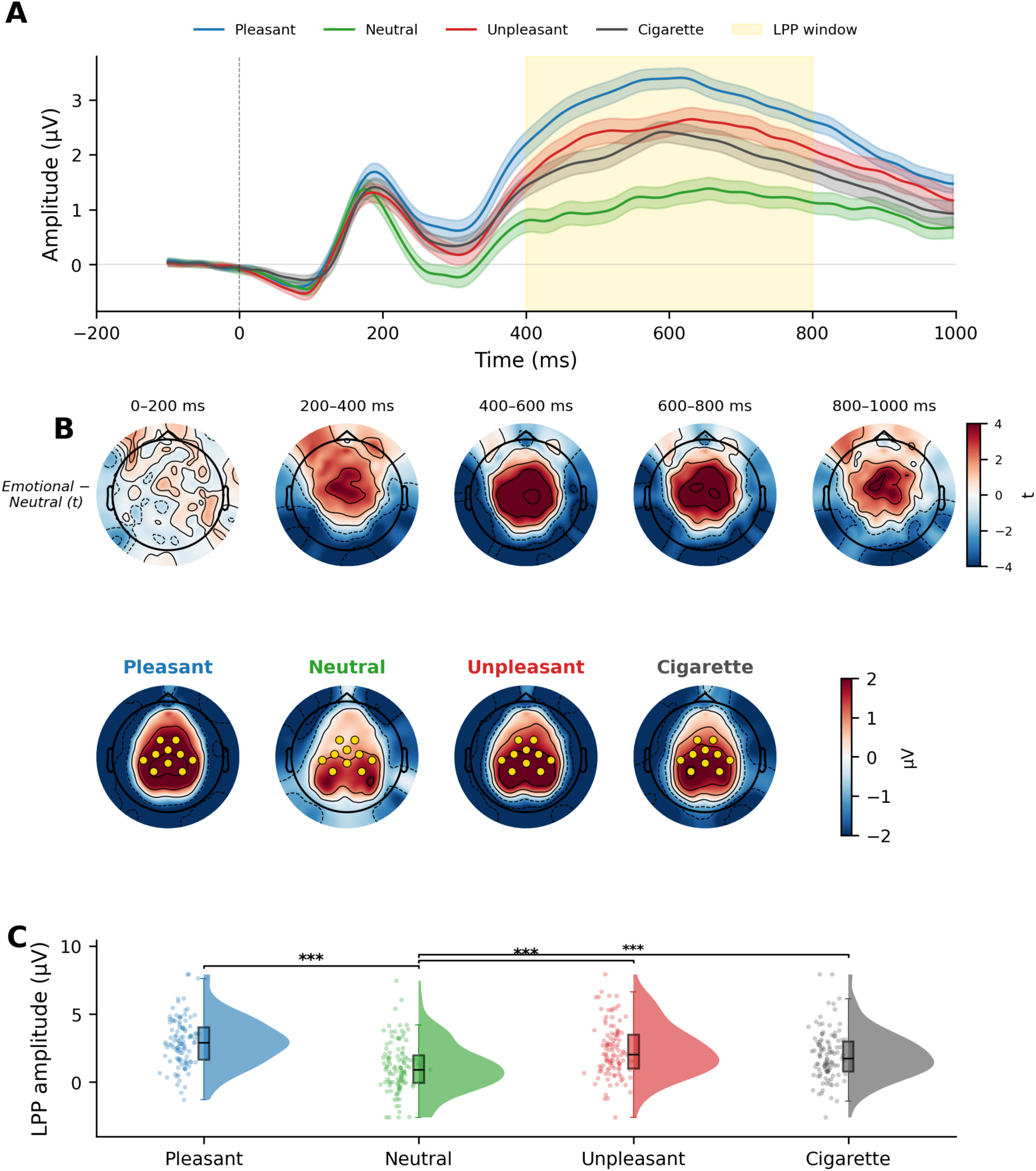
Emotional pictures elicit larger late positive potentials during the first (pre-instruction) picture presentation. (**A**) Grand-average ERP waveforms at centro-parietal ROI channels for pleasant (blue), neutral (green), unpleasant (red), and cigarette (gray) pictures during the first passive viewing. Shaded ribbons represent ±1 SEM. Yellow shading indicates the LPP measurement window (400–800 ms). (**B**) Top row: topographic maps of the Emotional − Neutral difference (*t*-statistic) across five 200-ms time bins, showing the emergence of the centro-parietal emotion effect. Bottom row: topographic maps of mean scalp voltage (µV) during the LPP window (400–800 ms) for each valence category. Gold dots indicate a priori ROI channels. (**C**) Raincloud plots of individual LPP amplitudes by valence category. All three stimulus categories elicited significantly larger LPPs than Neutral (all *p*s < .001). ***p < .001.

### Arousal modulates the cognitive load effect

Given the absence of any direction-specific instruction effect in the primary analysis (Enhance vs. Suppress BF01 = 9.30), the following analyses collapse Enhance and Suppress into a single Regulate condition to examine whether the overall attentional load effect varies with stimulus arousal. As a confirmatory check, Enhance and Suppress were also examined separately; the two conditions did not differ at either arousal level for either valence category (all BF01s > 3.5), justifying the collapsed approach. To test whether the cognitive load effect varied with stimulus intensity, we exploited the within-design arousal manipulation by comparing high-arousal and low-arousal stimuli separately. In the View condition (Supplementary Fig. 4A), pictures pre-selected to be included in the high-arousal category elicited substantially larger LPPs than low-arousal pictures, which in turn elicited larger LPPs than Neutral (all ps < .05). We then compared View to a collapsed Regulate condition (average of Enhance and Suppress) at each arousal level (Supplementary Fig. 4B). For high-arousal pictures, View produced significantly larger LPPs than Regulate for both pleasant and unpleasant stimuli (both ps < .05). For low-arousal pictures, the difference was small and nonsignificant. Critically, this outcome is inconsistent with the predictions of both the strong-situation and strategy-selection frameworks. Both frameworks predict that instruction-specific LPP modulation, that is, the Enhance > View > Suppress ordering, should be most evident at low arousal, when participants can fully implement the reappraisal strategy and volitional control is least constrained by stimulus intensity. Yet no instruction effect in any direction emerged at low arousal. The cognitive load pattern appeared exclusively at high arousal, where the emotional LPP is large enough to make attentional competition detectable.

Bayesian analyses revealed a complementary dissociation (Supplementary Fig. 4C). The Reappraisal contrast (Enhance − Suppress) yielded inconclusive-to-null evidence at both arousal levels, confirming that regulation direction did not modulate the LPP regardless of stimulus intensity. The View vs. Regulate contrast showed strong evidence for the alternative at high arousal (View > Regulate) but shifted toward the null at low arousal.

**Supplementary Figure 4.**
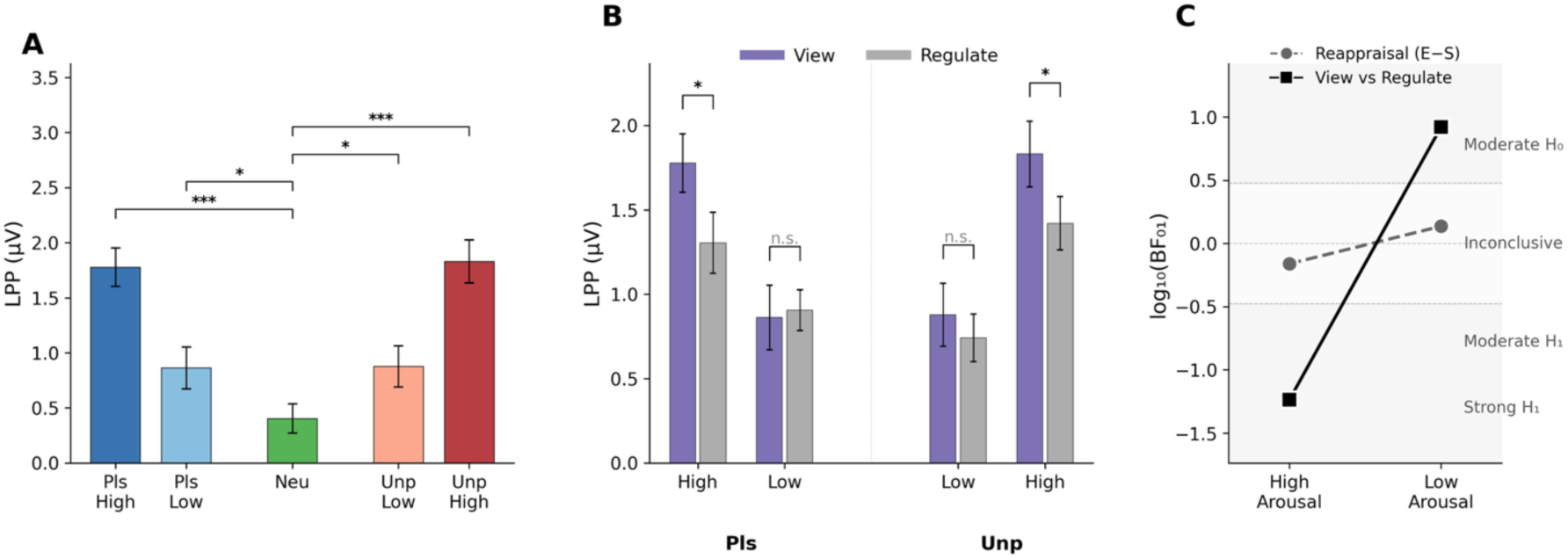
The cognitive load effect on the LPP is arousal-dependent. (A) Mean LPP amplitude (±SEM) during passive viewing for high- and low-arousal pictures. (B) Instruction effects (View vs. Regulate) for each arousal × valence cell. (C) Bayesian evidence slope graph.

## Cigarette Pictures

### Behavioral ratings

In the View condition, cigarette pictures were rated as more emotionally intense than neutral pictures, t(106) = 4.98, p < .001, d = 0.48, but lower than both pleasant, t(106) = −2.78, p = .006, d = −0.27, and unpleasant pictures, t(106) = −6.06, p < .001, d = −0.59 (Supplementary Fig. 5A). Regulation instructions modulated cigarette ratings in the same direction as emotional pictures. A one-way repeated-measures ANOVA showed a significant effect of Instruction, F(2, 212) = 48.72, p < .001, ηG² = .043, ε = .75. Enhance instructions increased ratings relative to View, t(106) = 9.40, p < .001, d = 0.91, and Suppress instructions decreased them, t(106) = −3.10, p = .003, d = −0.30 (Supplementary Fig. 5B). The Enhance–Suppress difference was large, t(106) = 7.92, p < .001, d = 0.77. These behavioral effects parallel the pattern observed for pleasant and unpleasant pictures in the main analysis, confirming that participants applied the regulation instructions consistently across all stimulus categories.

**Supplementary Figure 5.**
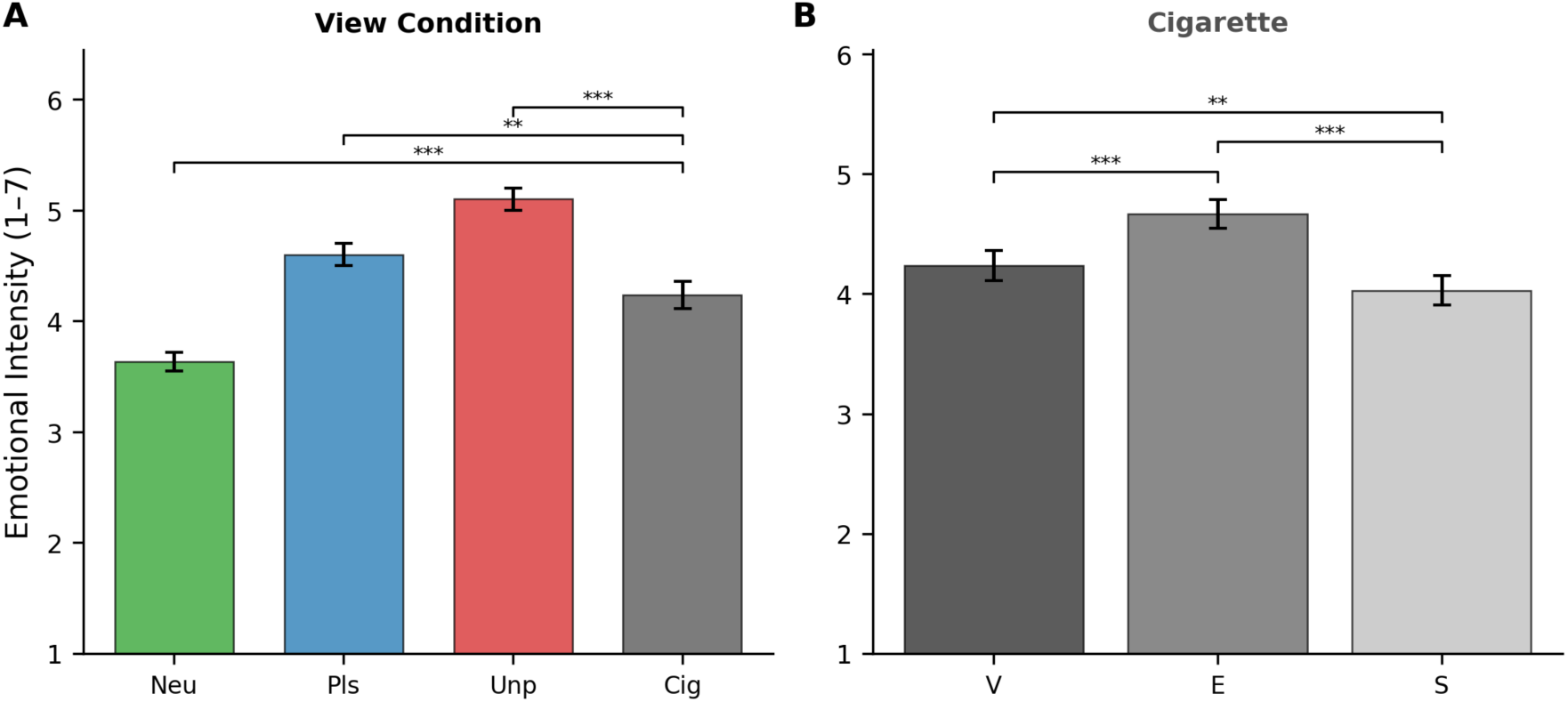
Behavioral ratings for cigarette-related pictures. (A) Mean emotional intensity ratings in the View condition across all four stimulus categories. Cigarette pictures were rated as more intense than neutral (p < .001, d = 0.48) but less intense than both pleasant (p = .006, d = −0.27) and unpleasant pictures (p < .001, d = −0.59). (B) Mean ratings for cigarette pictures under View, Enhance, and Suppress instructions. The instruction effect paralleled the pattern observed for emotional pictures, with Enhance increasing and Suppress decreasing ratings relative to View. Error bars represent ±1 SEM. **p < .01, ***p < .001.

### Late Positive Potential in the second presentation

In the pooled sample, cigarette pictures viewed passively (*M* = 0.46, *SD* = 1.65 µV) did not differ from neutral pictures (*M* = 0.37, *SD* = 1.43 µV), *t*(106) = 0.54, *p* = .588, *d* = 0.05, 95% CI [−0.14, 0.24] (Supplementary Fig. 6A). This null effect was consistent across samples: Sample 1, *t*(72) = 0.73, *p* = .465, *d* = 0.09; Sample 2, *t*(33) = −0.12, *p* = .907, *d* = −0.02. Grand-average waveforms confirmed that the cigarette time course tracked the neutral condition throughout the LPP window (400–800 ms), in contrast to the clear divergence of pleasant and unpleasant waveforms from neutral beginning around 300 ms (Supplementary Figure 6A).

Reappraisal instructions did not modulate LPP amplitude to cigarette pictures (Supplementary Fig. 6B). In the pooled sample, descriptive statistics were: View (*M* = 0.46, *SD* = 1.65 µV), Enhance (*M* = 0.60, *SD* = 1.33 µV), and Suppress (*M* = 0.49, *SD* = 1.58 µV). Pairwise comparisons were uniformly nonsignificant: View vs. Enhance, *t*(106) = −0.92, *p* = .358, *d* = −0.09 [−0.28, 0.10]; View vs. Suppress, *t*(106) = −0.21, *p* = .835, *d* = −0.02 [−0.21, 0.17] (Supplementary Figure 3B). This pattern was consistent in both cohorts: in the Discovery sample (*N* = 73), View vs. Enhance: *t*(72) = −1.23, *p* = .222, *d* = −0.15; View vs. Suppress: *t*(72) = −0.43, *p* = .667, *d* = −0.05; in the Replication sample (*N* = 34), View vs. Enhance: *t*(33) = −0.01, *p* = .994; View vs. Suppress: *t*(33) = 0.34, *p* = .737.

Bayesian model comparison confirmed the absence of any instruction effect for cigarette pictures (Supplementary Fig. 6C). The Reappraisal contrast (Enhance − Suppress) yielded BF01 = 7.5, indicating moderate evidence for the null. The Cognitive Load contrast (2 × View − Enhance − Suppress) yielded BF01 = 7.5, similarly favoring the null. The combined Null model received very strong support (BF01 = 56.1). This pattern was consistent across samples (Discovery: combined null BF01 = 32.3; Replication: BF01 = 28.4). The absence of both emotion and instruction effects for cigarette pictures is internally consistent. Because cigarette images did not engage the motivational systems indexed by the LPP, there was no emotional response to modulate with reappraisal instructions, nor differential attentional capture to reduce cognitive load. This null pattern serves as an informative negative control, demonstrating that the instruction effects observed for pleasant and unpleasant stimuli were not an artifact of task demand or nonspecific compliance.

Since visual inspection of the grand-average waveforms (Supplementary Figure 3A) suggested an early modulatory effect for cigarette pictures relative to neutral, we conducted a sub-window analysis, splitting the LPP epoch into early (400–600 ms) and late (600–800 ms) halves. The Cigarette–Neutral difference was nonsignificant in both sub-windows (400–600 ms: *t*(106) = 1.77, *p* = .080, *d* = 0.17, BF₀₁ = 5.6; 600–800 ms: *t*(106) = 0.13, *p* = .898, *d* = 0.01, BF₀₁ = 25.8), whereas Pleasant and Unpleasant pictures remained significantly above Neutral in both sub-windows (all *p*s < .001, *d*s = 0.48–0.67). The apparent early divergence thus reflects a transient, unreliable deflection rather than sustained motivational engagement, as indexed by the LPP.

**Supplementary Figure 6.**
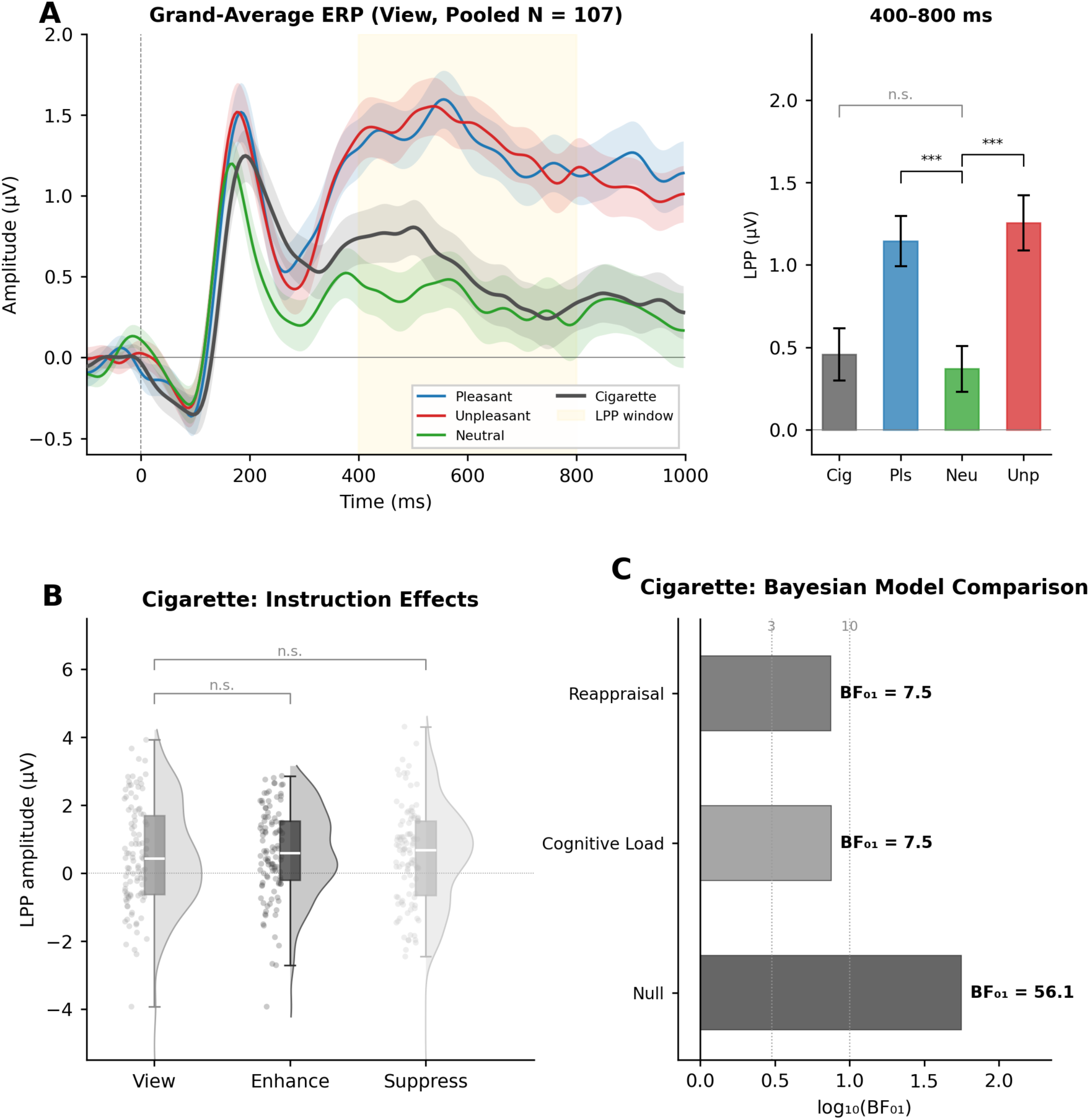
Cigarette-related stimuli: LPP results. **(A)** Grand-average ERP waveforms at centro-parietal ROI channels in the View condition for all four stimulus categories (left) and mean LPP amplitude (400–800 ms) with ±1 SEM error bars (right). Cigarette pictures did not differ from the Neutral condition. \*\*\**p* < .001, n.s. = not significant. **(B)** Raincloud plots of LPP amplitude for cigarette pictures under View, Enhance, and Suppress instructions. No pairwise comparisons reached significance. **(C)** Bayesian model comparison for cigarette pictures. All three models favor the null hypothesis, with the combined Null model receiving very strong support (BF01 = 56.1). Dotted lines indicate BF = 3 and BF = 10 evidence thresholds.

## Notes

https://github.com/nsambuco/LPP_reappraisal

### Competing Interest Statement

The authors have declared no competing interest.

